# Ejaculate deterioration with male age, and its amelioration in *Drosophila*

**DOI:** 10.1101/624734

**Authors:** Irem Sepil, Ben R Hopkins, Rebecca Dean, Eleanor Bath, Solomon Friedman, Ben Swanson, Harrison J Ostridge, Norene A Buehner, Mariana F Wolfner, Rebecca Konietzny, Marie-Laëtitia Thézénas, Elizabeth Sandham, Philip D Charles, Roman Fischer, Josefa Steinhauer, Benedikt M Kessler, Stuart Wigby

## Abstract

Declining ejaculate performance with male age is taxonomically widespread and has broad ramifications for fertility and fitness. However, we have a poor understanding of age-related changes to specific ejaculate components, how they cause reduced performance, and whether the decline is ameliorable. Here, we show that, in *Drosophila*, sperm production chronologically declines with age, invariant to mating activity, while repeated mating causes infertility via reduced sperm stores and viability. However, changes to sperm do not fully explain ejaculate deterioration: impacts on seminal fluid contribute to aspects of reduced ejaculate performance, associated with shifts in proteome abundance and quality. We show that ablation of insulin-like peptide-producing cells in males ameliorates aspects of ejaculate performance loss, suggesting that anti-ageing interventions can be coopted to benefit male reproductive health.

**One Sentence Summary:** Ejaculate performance declines with male age via mating-dependent sperm and seminal protein deterioration, but it can be ameliorated.

## Main Text

There is accumulating evidence that increased male age reduces ejaculate performance across a wide range of animal taxa (*1, 2*). Age-related declines in ejaculate performance have broad ramifications for reproductive biology and sexual selection, and in humans they contribute to the current “male fertility crisis” (*3*) due to trends for delayed fatherhood (*4*). However, the mechanisms underlying age-related declines in male ejaculate performance are poorly understood (*1*) and – despite major recent advances in the biology of ageing (*5*) – it remains unclear whether they are ameliorable. We typically do not know how male age impacts specific components of the ejaculate and crucially, how these changes link to fertility and other aspects of ejaculate function, such as stimulating post-mating changes to female physiology and behaviour. Much past research in the field has focussed on ageing impacts on sperm or testes (*6*–*8*), but the ejaculate is composed of both sperm and non-sperm seminal fluid. The seminal fluid is a complex cocktail of functionally diverse molecules that makes crucial contributions to ejaculate function (*9, 10*). The seminal fluid proteins (Sfps) in particular are central in supporting sperm function, and in many species also modulate post-mating female physiology and behaviour (*11*). *Drosophila melanogaster* is a well-established model for both ageing and ejaculate research, but these fields have largely operated independently (but see *12*–*15*). Here, we use *D. melanogaster* to identify mechanisms that both drive and slow the loss of ejaculate performance with male age. We dissect the contributions of age-related changes to sperm and Sfps to a suite of key ejaculate functions in this species (e.g. fertility, fecundity, sperm competitiveness, female refractoriness), and investigate the impact of a somatic lifespan-extending intervention on these functions.

### Reproductive consequences of male ageing and mating history

We measured reproductive traits in experimental males that were one-week-old (1w), three-weeks-old (3w) or five-weeks-old (5w) and had been maintained in either single-sex groups of twelve (unmated, “U”), or in groups of three males and nine females (frequently mated, “F”) (Fig. 1A). These time points span male peak reproductive performance (1w, young) and reproductively senesced states (5w, old) while ensuring that most males survive the experiment (*14*) (fig. S1). As expected based on previous work (*12, 15, 16*), we found clear evidence that male reproductive function declines with age. However, the effects are highly dependent on male sexual activity. Old-F males father fewer offspring, are more likely to be infertile, are poorer at suppressing female remating and their sperm perform poorly when competing with the ejaculates of rival males (Fig. 1B-E). Old, unmated males (Old-U) are also poor sperm competitors, but their reproductive output, fertility and ability to suppress female remating are not significantly reduced compared to young males (Fig. 1B-E). Old males also show a significant reduction in copulation probability – an effect again exacerbated by frequent mating (fig. S2). Taken together, our results highlight how frequent mating activity is an important contributor to age-dependent reproductive decline in males. Reduced copulation probability likely results from the reduced courtship ability of old males and female discrimination of Old-F mated males (*17*), while the decline in post-mating traits signifies male age impacts on the ejaculate.

**Fig. 1:**
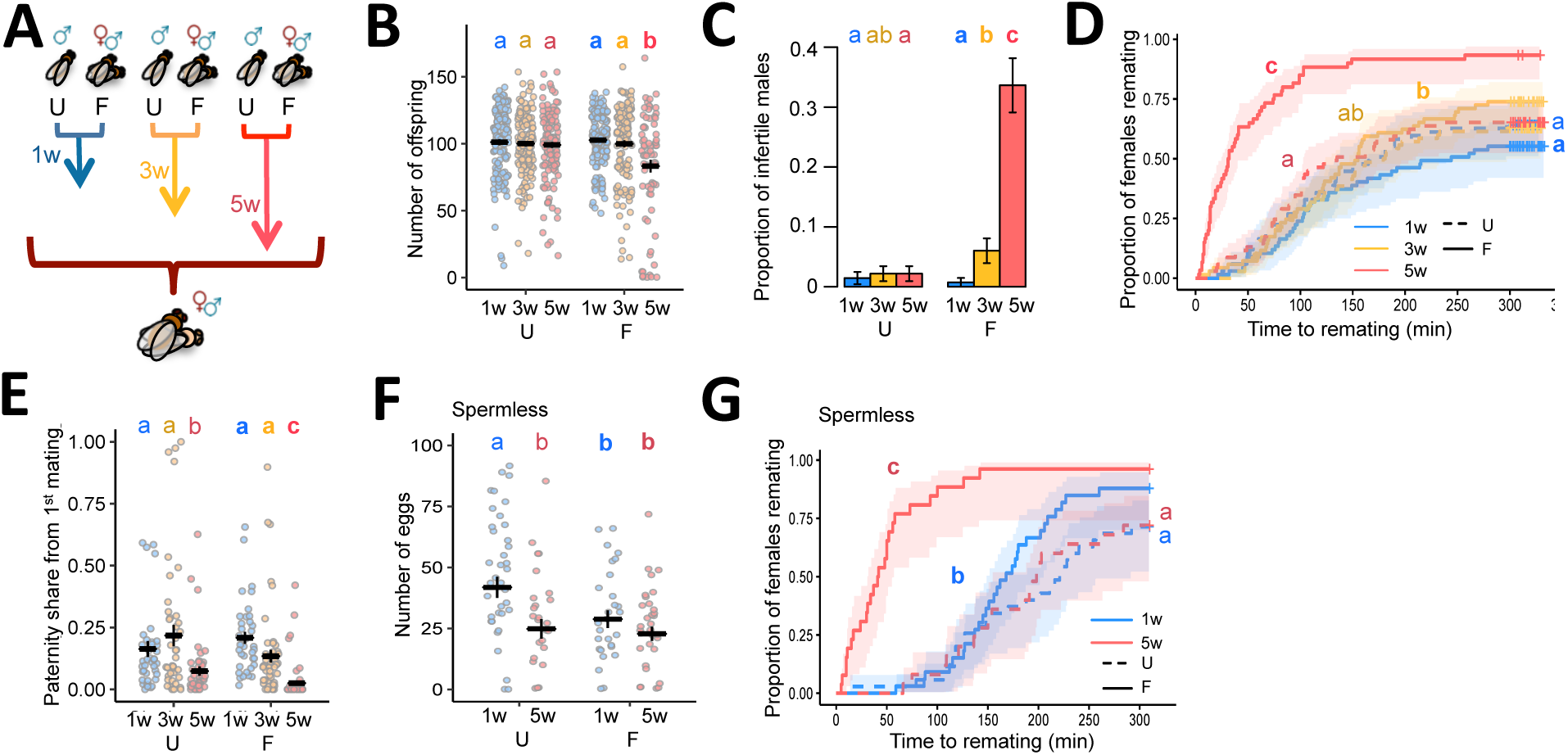
Decline in reproductive performance in response to male age and mating activity. **(A)** Experimental design. Unmated (U) and frequently mated (F) males. **(B)** Number of offspring from a single mating, excluding infertile matings (age and mating interaction: χ ^2^_2_ = 98.668; p= 0.0005) (n= 73-135). **(C)** The proportion of infertile matings (age and mating interaction: χ ^2^_2_ = 11.32; p= 0.0035) (n= 110-137). **(D)** Female latency to remate (age and mating interaction: χ ^2^_2_ = 34.886; p< 0.0001) (n= 60-70). **(E)** Paternity share of the experimental first male (age and mating interaction: χ ^2^_2_ = 219.34; p= 0.0234) (n= 37-55). **(F)** Number of eggs laid by females mated to spermless experimental males (*binomial*: age: χ ^2^_1_ = 1.2509; p= 0.263; mating: χ ^2^ = 0.609; p= 0.435; age and mating interaction: χ ^2^_1_ = 0.027; p=0.87; *count:* age: χ ^2^_1_ = 83.106; p= 0.003; mating: χ ^2^_1_ = 102.38; p= 0.0008; age and mating interaction: χ ^2^_1_ = 12.774; p= 0.237) (n= 30-42). **(G)** Female latency to remate (age and mating interaction: χ ^2^_1_ = 13.648; p= 0.0002) (n= 25-35). Results shown as means ± SEM. Shaded areas confidence intervals at 0.15 level. Differences at p < 0.05 within mating groups and age categories represented as different letters.

Next, we determined whether the seminal fluid alone, in the absence of sperm, contributes to age-related reproductive decline. Long-term elevated egg production and sexual refractoriness in females requires the receipt of sperm as well as Sfps, but these responses can also be partially elevated in the short-term without sperm (*18*). Using spermless (*son-of-tudor*) males we found that old males and young, frequently mated (Young-F) males are poorer at stimulating female fecundity (Fig. 1F and fig. S3). Likewise, Old-F males and, to a lesser extent, Young-F males are poorer at suppressing female remating (Fig. 1G and fig. S3). As the ejaculates of *son-of-tudor* males do not contain sperm, the age-related decline in reproductive function in these experiments is due to Sfps. This is because Sfps are known to be the seminal fluid component that stimulates fecundity and refractoriness responses in females (*11*).

### Age and mating effects on the seminal proteome

Having identified loss of ejaculate performance associated with increased age and mating activity, and the direct contribution of the seminal fluid, we next investigated changes to the seminal fluid proteome to explain these effects. We first applied label-free quantitative proteome analysis to the Sfp-producing tissues (accessory glands and ejaculatory duct) of experimental males (*19*).

We focused our analyses on established Sfps (*19, 20*) and first examined their production by males before transfer to females. We found that the abundance of many Sfps increases with age in unmated males, mirrored by an increase in size of the accessory gland, the tissue that makes most Sfps, whereas Sfp abundance and accessory gland size do not change with age in frequently mated males (Fig. 2 and fig. S4). A principal component analysis supported these findings, showing that the composition of the seminal fluid proteome changes significantly with age in unmated males, but not in frequently mated males (Fig. 2C). All of the upregulated Sfps are specific to the accessory glands and include Sfps that function in sperm storage and female post-mating behavior modification (*11*) (table S1). In contrast, the abundance of most of the ejaculatory duct derived Sfps (*19, 21*) do not exhibit a differential response to age and mating; they cluster separately from the rest of the proteins (Fig. 2A and table S2). Notably, this disparity indicates that the two male reproductive tissues respond differentially to age and mating.

**Fig. 2:**
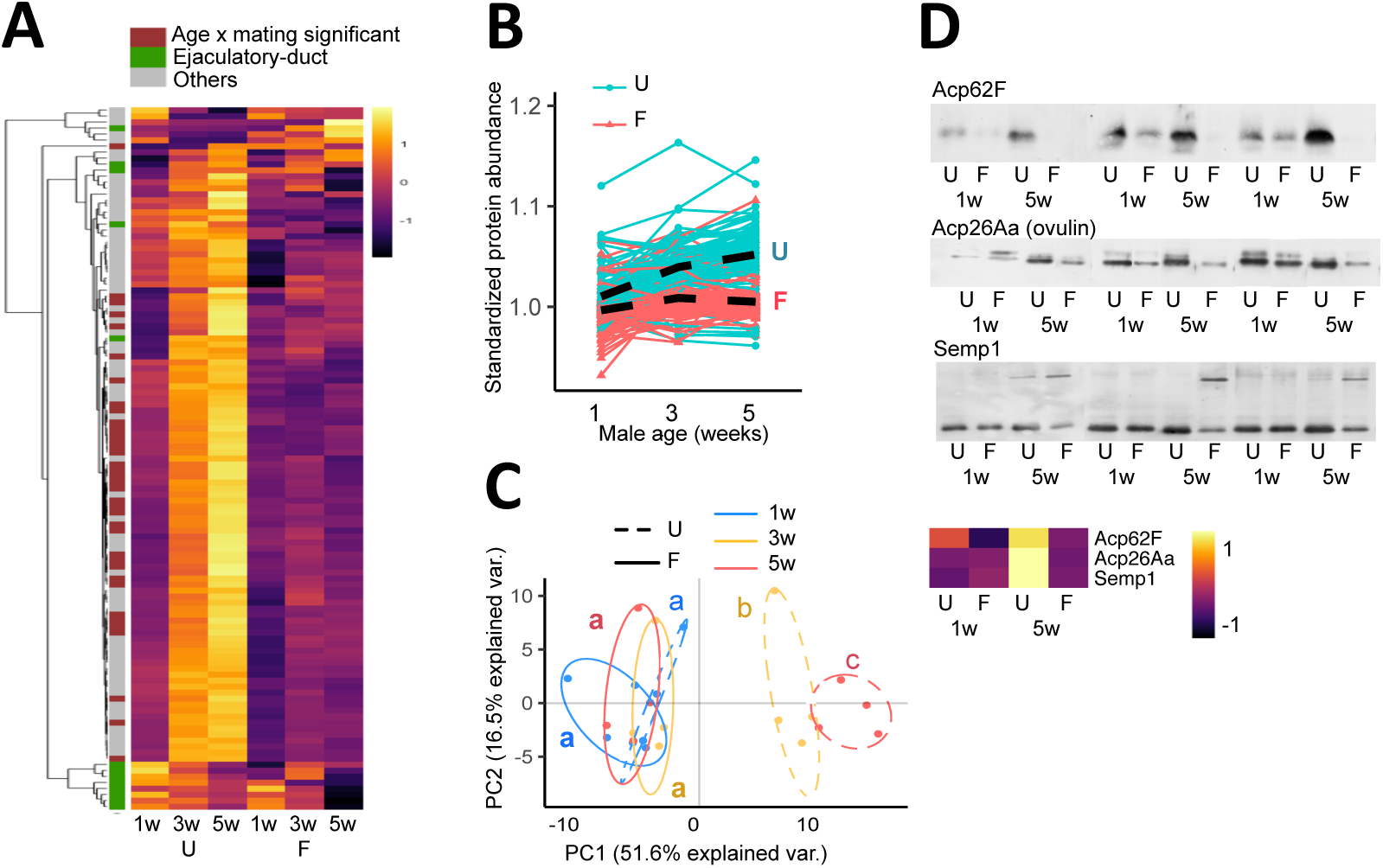
The seminal fluid proteome responds differentially to ageing in U versus F males. **(A)** Heatmap of the abundance of the 117 Sfps detected in accessory gland and ejaculatory duct samples (n=4 replicate experiments per group). The abundance of 40 out of 117 Sfps exhibit a significant differential response to age and mating after multiple test correction. The annotation classification of each Sfp is indicated. **(B)** Line plots showing the change in standardized Sfp abundance with age. The average change in Sfp abundance for unmated and frequently mated males is depicted with lines marked ‘U’ and ‘F’ respectively (age and mating interaction: L Ratio^2^_2_ = 163.856; p< 0.0001). **(C)** Principal component analyses of the seminal fluid proteome in male reproductive tissues (age and mating interaction: L Ratio^2^_2_ = 34.949; p< 0.0001). **(D)** Representative Acp62F, Acp26Aa and Semp1 Western blots, in 1w and 5w old unmated (U) and frequently mated (F) males. The abundance of each protein is predicted from the proteomic data and illustrated as a heatmap. Each lane is an individual male. Full blots are shown in fig. S5.

Next, we performed Western blot analyses for six Sfps of known functional importance to look for evidence of qualitative changes related to age and mating history. For each Sfp tested, our proteomic data, which are based on trypsin-cleaved peptides, indicate either an age-related increase in unmated males or no change in frequently mated males in Sfp abundances. However, Acp62F, an Sfp which has been implicated in sperm competition (*22*), is largely undetectable by Western blot in Old-F males (Fig. 2D and fig. S5). This suggests that ageing degrades Acp62F in such a way that while no band is detectable on Western blot, the trypsin-cleaved peptides remain identifiable by mass spectrometry. The double band of Acp26Aa (ovulin), one of the most rapidly evolving proteins in *Drosophila* (*23*), becomes either more condensed or loses the top band altogether in old males, representing possible age-specific alternative splicing or post-translational modification. Likewise, Semp1, a seminal metalloprotease that cleaves ovulin within females (*24*), shows an additional upper band in old males from both mating groups, indicating a potential age-specific post-translational modification or attachment to a larger protein (Fig. 2D and fig. S5). Full-length ovulin and two of its cleavage products stimulate ovulation, hence any deterioration with age and mating would negatively impact female ovulation rate following mating (*25*). We saw no qualitative changes in the three other Sfps tested (Acp70A [sex peptide], Acp36DE and CG9997; fig. S5). Together, these results indicate that a subset of seminal proteins display qualitative alterations in response to age and, in some cases the combination of age and frequent mating. These qualitative changes are consistent with a loss of seminal protein homeostasis (*26*) and are associated with the compromised post-mating phenotypes in females mating with old males.

By comparing the quantity of Sfps present in males before and after mating we can infer Sfp transfer to females during copulation (*19*). We found that the abundance of Sfps transferred changes significantly with age and mating, leading to a significant change in Sfp proteome composition. Transferred Sfps show an age-related decline in unmated males (Fig. 3), in spite of their higher accumulation in the accessory glands (Fig. 2): i.e. despite producing Sfps in greater abundance, old-U males appear to be poor at transferring them to females during copulation. We again observed separate clustering for several ejaculatory duct specific Sfps, where the trend is an increase in Sfp transfer with age in frequently mated males, although this effect was weaker than the differences seen in Sfp production. Together these results suggest that a distinct set of Sfps accumulates with age in the absence of mating, resulting in compositional change of the seminal proteome. In contrast, the abundance and transfer of Sfps is maintained with age in the presence of frequent mating, which, given that their striking decline in ejaculate performance, indicates that quantitative Sfp effects do not cause male age-related ejaculate deterioration in these sexually active males.

**Fig. 3:**
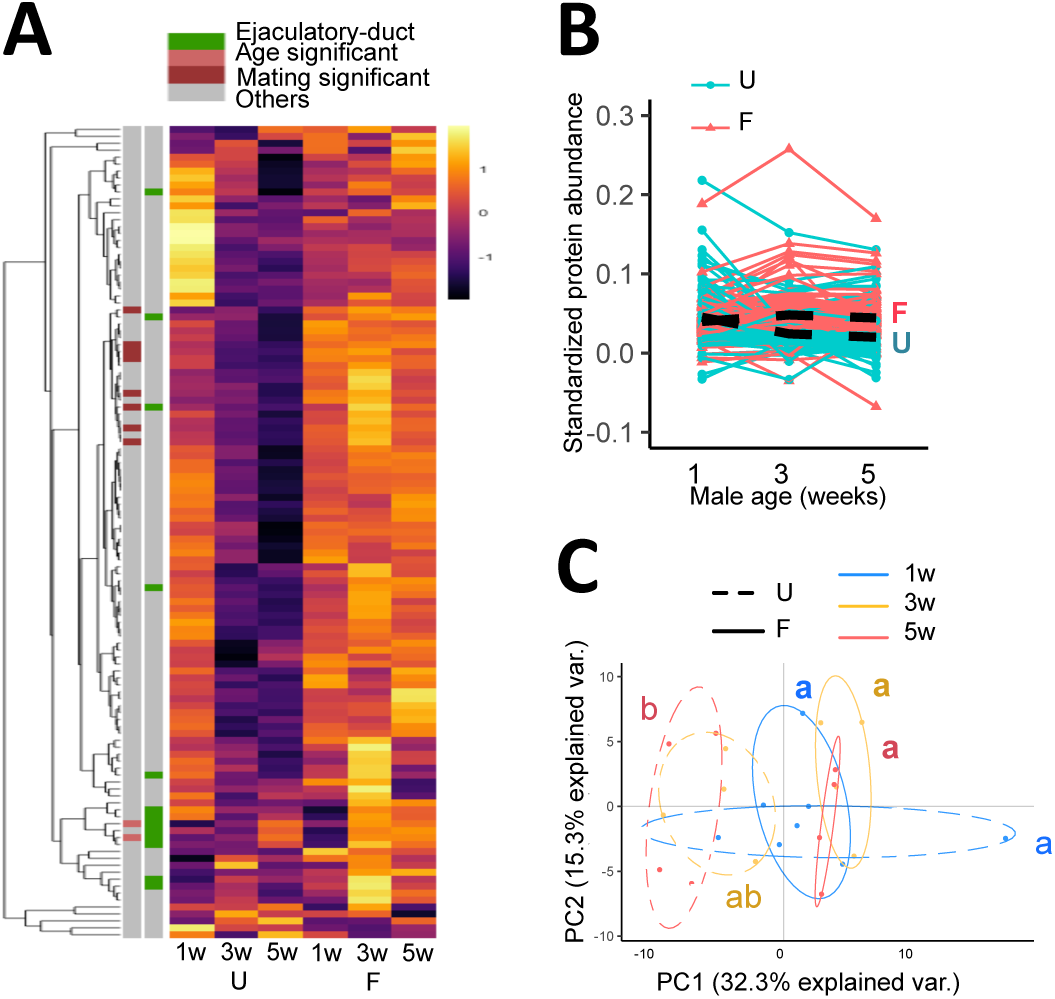
The seminal fluid proteome transferred to females responds differentially to ageing in U versus F males. **(A)** Heatmap of the abundance of 117 seminal fluid proteins transferred to females during mating. None of the individual 117 Sfps exhibited a significant interaction between age and mating group after false discovery rate multiple test correction. Two ejaculatory duct-specific Sfps were transferred in significantly higher quantities in response to age, independent of mating activity (CG17242, CG5162), and ten Sfps were transferred in significantly higher quantities in response to frequent mating independent of age (Acp26Aa, CG10587, CG17472, CG3097, CG34002, Est-6, NLaz, Regucalcin, Sfp24F, Sfp65A). The annotation classification of each Sfp is indicated. **(B)** Line plots showing the standardized abundance of Sfps transferred with age. The average change in Sfp abundance for unmated and frequently mated males is depicted with lines marked ‘U’ and ‘F’ respectively (age and mating interaction: L Ratio^2^_2_ = 130.595; p< 0.0001). **(C)** Principal component analyses of the seminal fluid proteome transferred to females (age and mating interaction: L Ratio^2^_2_ = 11.485; p= 0.003). Differences at p < 0.05 within mating groups and age categories are represented as different letters.

Finally, we investigated whether the previously-identified sperm-protecting function of seminal fluid (*27*) declines with age and frequent mating. Using SYBR-14 and propidium iodide fluorescent staining, we measured the effects of seminal fluid on the survival of sperm recovered from a different male. However, we found no evidence that age or mating history compromises the ability of seminal fluid to keep sperm alive (fig. S6).

### Age and mating effects on sperm

Consistent with previous evidence of declining rates of spermatogenesis with age in flies (*28*), we found that the number of germline cysts in the final individualization stage of spermatogenesis (*29*) declines substantially as males age. Strikingly, this decline occurs at indistinguishable rates in unmated and frequently mated males. This suggests that males undergo a chronological decline in sperm production which is invariant to mating activity (Fig. 4A). This finding is surprising given that sperm production rate is known to be malleable; for example, males elevate sperm production in response to the presence of rivals (*30*), and testis germline stem cell maintenance responds plastically to nutrition (*31*).

**Fig. 4:**
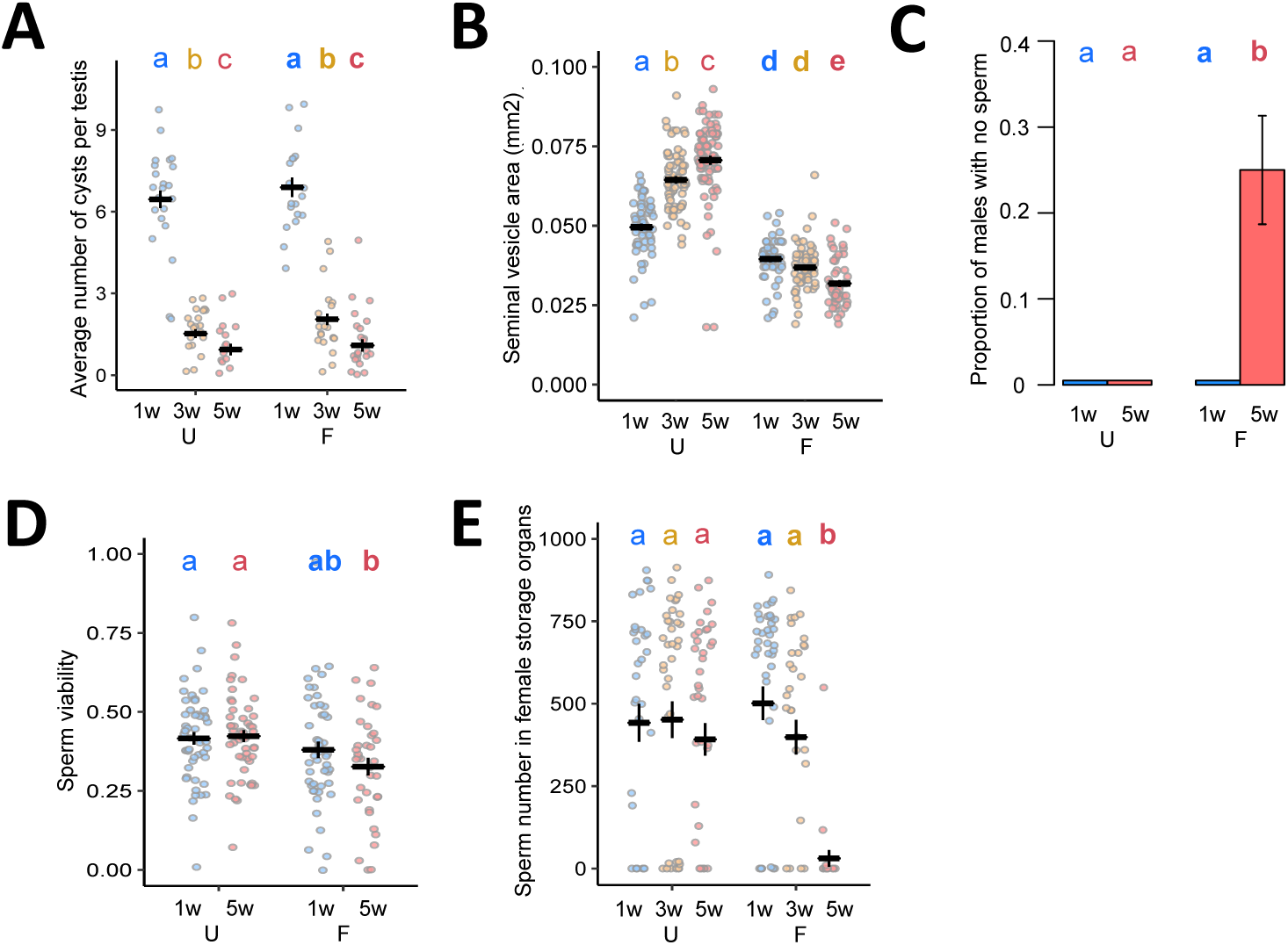
Ageing and frequent mating impacts sperm production and transfer. **(A)** Average number of mature germline cysts per testis (a measure of sperm individualization rate) (age: χ ^2^_2_= 223.78; p< 0.0001; mating: χ ^2^_1_= 1.334; p= 0.119; age and mating interaction: χ ^2^_2_= 0.772; p= 0.496) (n= 16-22). **(B)** Seminal vesicle area (mm^2^) (age and mating interaction: F^2^_2_ = 68.494; p< 0.0001) (n= 49-79). **(C)** Proportion of males with no evidence of sperm within the seminal vesicle (age and mating interaction: χ ^2^_1_ = 1; p< 0.0001) (n= 47-51). **(D)** Sperm viability 10 minutes after removal from the seminal vesicles (male age: χ ^2^_1_ = 0.004; p= 0.985; mating group: χ ^2^ = 102.69; p= 0.002; interaction between male age and mating group: χ ^2^_1_ = 15.921; p= 0.223) (n= 36 - 51). **(E)** Number of GFP fluorescent sperm heads in female sperm storage organs 90 minutes after mating starts (*binomial*: age and mating interaction: χ ^2^_2_ = 13.417; p= 0.0012; *count*: age and mating interaction: χ ^2^_2_ = 726.46; p= 0.0062) (n= 22-43). Results are shown as means ± SEM. Differences at p < 0.05 within mating groups and age categories are represented as different letters. “U” stands for unmated and “F” stands for frequently mated males.

In the absence of mating, the size of the seminal vesicles (where mature sperm are stored) increases in unmated males, but decreases in frequently mated males (Fig. 3B). This suggests that, like Sfps in the accessory glands, sperm stores accumulate in unmated males, despite the declining rate of sperm production. However, in contrast to Sfps, males are unable to sufficiently replenish sperm when they mate frequently throughout life, leading to depletion of sperm stores and high incidences of no sperm being found in the seminal vesicles of Old-F males (Fig. 3C). Frequently mated males also have lower sperm viability, independent of age class, suggesting that regular copulation leads to reduced sperm quality (Fig. 3D). As might then be expected, we found a significant reduction in the number of sperm present in sperm storage organs (seminal receptacle and spermatheca) of females mated to Old-F males, relative to all other treatments, although there were also nonsignificant downward trends for both Old-U and 3 week-F males (Fig. 3E).

The fact that Old-F males are sperm-depleted, but show no evidence of decline in Sfp quantity, indicates that a mismatch develops in the relative capacity to produce sperm and Sfps. In the short term, when males mate several times in rapid succession, seminal fluid rather than sperm is thought to limit fertility in male *Drosophila* and other insects (*32*), but our data show that over the long term, Sfp replenishment capacity remains strong and is little affected by age. However, our data clearly show that non-sperm components of the ejaculate also contribute to the loss of ejaculate performance in ageing males, as evidenced by a reduced ability to stimulate female post-mating responses (Fig. 5), known to be controlled by Sfps (*11*). Old-U males, despite accumulating Sfps with age, transferred reduced Sfp amounts to females, which could contribute to their decline in ejaculate performance, while Old-F males did not show a reduction in Sfp transfer, yet they had poorer reproductive performance. It is therefore likely that simplistic quantitative changes in Sfps play at best a minor role in declining ejaculate performance. Instead, our data are consistent with the idea that qualitative seminal fluid defects, such as a loss of seminal protein homeostasis, and compositional changes in unmated males, contribute to age-related declines in ejaculate performance. These seminal fluid defects occur in concert with declines in sperm numbers and viability in Old-F males (Fig. 5).

**Fig. 5:**
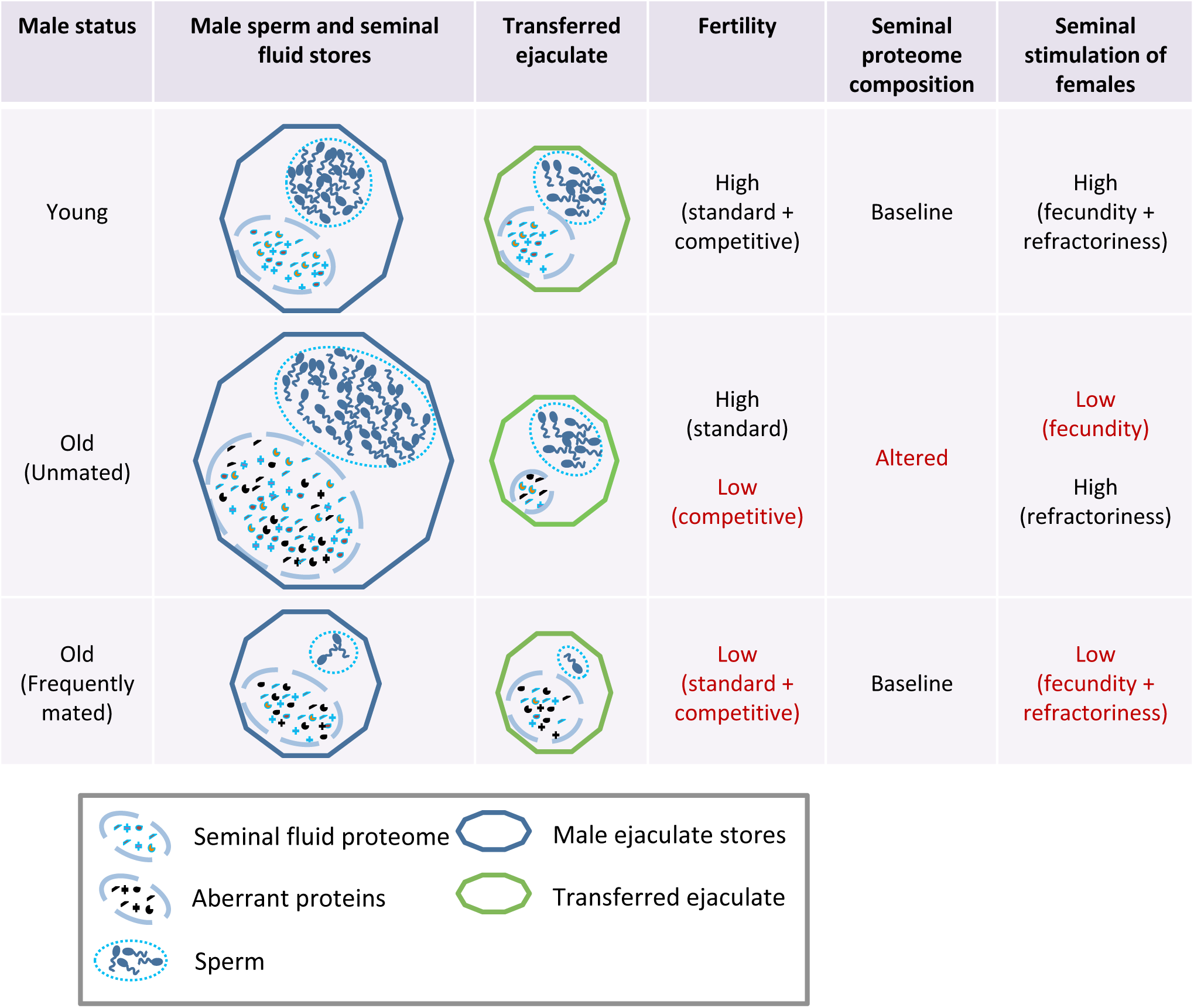
Schematic summary of the impacts of ageing on the ejaculate.

### Changes in reproductive ageing with lifespan extension

Manipulations of the insulin-signaling pathway can extend lifespan in a broad range of taxa (*33, 34*), but it is unclear if lifespan extension results in a trade-off with male reproductive function, or whether it could provide co-benefits to late-life ejaculate health. We used males in which the insulin-like peptide (dilp)-producing median neurosecretory cells (mNSCs) were ablated late in development (*35*) (hereon ‘ablated males’) and confirmed that these males display increased survival (fig. S7). We found clear evidence that the ablated males have reduced reproductive senescence. Old-F ablated males are significantly less likely to be infertile and significantly better at suppressing female remating than Old-F control males (Fig. 6). We did not detect any significant differences between ablated and control males in offspring production and paternity share (fig. S8). We also found that old ablated males are more likely to successfully copulate than controls, although the effect was more striking in the frequently mated treatment (fig. S9).

**Fig. 6:**
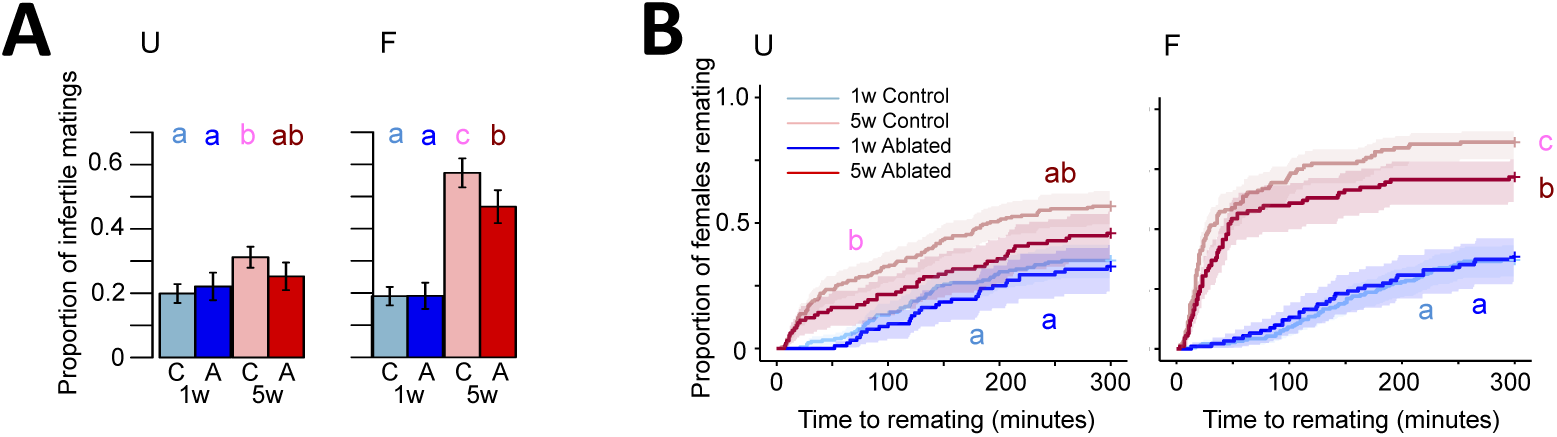
Manipulation of the insulin signaling pathway ameliorates ejaculate deterioration in Old-F males. **(A)** The proportion of infertile matings (*U:* age: χ ^2^_1_= 10.413; p= 0.001; line: χ ^2^_1_ = 0.437; p= 0.508; age and line interaction: χ ^2^_1_ = 1.941; p= 0.164; n= 95-202) (*F:* age: χ ^2^_1_ = 115.83; p< 0.0001; line: χ ^2^_1_ = 5.433; p= 0.0198; age and line interaction: χ ^2^_1_ = 3.684; p= 0.055; n= 94-189). (**B)** Female remating latency (*U:* age: χ ^2^_1_ = 28.927; p< 0.0001; line: χ ^2^_1_ = 3.602; p= 0.058; age and line interaction: χ ^2^_1_ = 0.660; p= 0.416; n= 92-198) (*F:* age and line interaction: χ ^2^_1_ = 4.727; p= 0.03; n= 91-186). “U” stands for unmated and “F” stands for frequently mated males. “C” stands for control and “A” stands for ablated lines. Results are shown as means ± SEM. Shaded areas are confidence intervals at 0.15 level. Differences at p < 0.05 within lines and age categories are represented as different letters.

Our data show that inhibition of the insulin signaling pathway, in addition to extending lifespan, can ameliorate at least some aspects of age-related loss of ejaculate performance. This result supports the idea that it is possible to simultaneously reduce both somatic and reproductive ageing in males. This result apparently contrasts with the effects of rapamycin in mice, which extends lifespan, but causes testicular degeneration (*36*). There are a number of possibilities that could explain the discrepancy, including differential action of insulin and rapamycin pathways in male reproductive organs as well as taxon-specific responses to nutrient-sensing pathway interventions.

## Conclusions

In humans, a number of studies have shown declines in sperm ejaculate volume, sperm count, motility and viability with age, patterns which are often – but not always – seen in other animals (reviewed in *37*). *Drosophila* show age-related changes in gene expression in some Sfps (*13*), while Red junglefowl show associations between distinct seminal proteome profiles and sperm speed in aging males (*38*). However, in general non-sperm components of the ejaculate have received little attention in the context of age-related declines in ejaculate performance. Our *Drosophila* study provides a uniquely comprehensive exposition of sperm and seminal proteome changes with age, and links these with a suite of fitness-related ejaculate performance phenotypes. Our data show that both the quantity and quality of sperm and seminal fluid proteins can contribute to the age-related decline in male ejaculate performance, but that the role of these different factors is highly-dependent on the mating environment. Moreover, our data indicate that organism-level insulin signaling is a mediator of both organism survival and male ejaculate quality retention, and represents a viable target for developing interventions to ameliorate the impact of ageing on the ejaculate.

Perhaps surprisingly, quantitative declines in seminal protein production seem to play a minimal role in age-related ejaculate deterioration in *Drosophila*, although seminal proteome imbalance and sub-optimal transfer may contribute in sexually abstinent males, perhaps due to harmful accumulation within the accessory glands that prevent normal ejaculation. However, the most striking fertility declines were strongly associated with a numerical loss of sperm resulting from a declining sperm production rate, which leads to a failure to replenish lost sperm stores. Sperm production declines are striking in humans: the daily rate approximately halves between the ages of 20 and 60 (*39*). A prime candidate for interventions to delay age-related ejaculate deterioration must therefore be to target the root of declining sperm production. Whether the improvements to late life fertility we observed in ablated males were a result of direct influences of reduced insulin activity in male reproductive tissues, or as part of overall organismal health (or both), remain to be elucidated. However, given that nutrient-sensing pathways are highly evolutionary conserved in their effects on lifespan and reproduction and that there is considerable overlap in the process of spermatogenesis, sperm proteomes, Sfp-producing cells, and the categories and function of Sfps (*10, 40, 41*), flies clearly represent a powerful system for developing the basic tools for healthy male reproductive ageing.

## Acknowledgements

This study is dedicated to the late Ian Moore, Fellow of Wadham College, who initiated our collaboration with the TDI. We thank Jennifer Perry, Juliano Morimoto and Jacob Callear for helping with the experiments.

## Author Contributions

I.S. and S.W. designed and supervised the study; I.S., B.R.H., E.B., S.F., B.S., H.O., N.A.B., M.F.W., R.K., M.-L.T., E.S., R.F., J.S., B.M.K. and S.W. performed research; I.S., R.D. and P.D.C. analysed the data; I.S. and S.W. wrote the paper with input from all the co-authors.

## Data and materials availability

LC-MS/MS data have been deposited to the ProteomeXchange Consortium via the PRIDE partner repository with the data set identifier PXD009451 (http://proteomecentral.proteomexchange.org). Other data, materials, associated protocols and technical details are available to researchers desiring to replicate or expand studies of *Drosophila* reproductive ageing.

## Supplementary Materials

Materials and Methods

Figs. S1 to S9

Tables S1 to S3

References (42-52)

