## Supplementary Materials for "Ejaculate deterioration with male age, and its amelioration in *Drosophila*"

**This PDF file includes:**

Materials and Methods

Figs. S1 to S9

Tables S1 to S3

**S1 Materials and Methods**

**S1.1 Stock and Fly Maintenance**

All flies were maintained at 25 °C on a 12:12 L:D cycle and fed Lewis medium (*42*). Adult flies were maintained in 36 mL plastic vials containing 7.5ml Lewis medium supplemented with *ad libitum* live yeast grains. Flies were reared using a standard larval density method by placing approximately 200 eggs on 50 mL of food in 250 mL bottles (*43*). Virgins were collected on ice anesthesia within 8h of eclosion and were randomly assigned to their experimental group. Unless stated otherwise, we used a lab-adapted, outbred *Dahomey* (*Dah*) wild-type stock for all our experimental males. This stock has been maintained in large population cages with overlapping generations since 1970.

To produce spermless males, we used *son-of-tudor* males (*44*). This was achieved by crossing homozygous *tudor* females with wild-type *Dahomey* males, and collecting the male progeny. These males have no germline, so do not produce sperm. Control males were of the same genetic background, but from non-homozygous mothers. To produce males that had fluorescently tagged sperm, we used *Ub-GFP* males (*45*). *Ub-GFP* males were created by germline transformation of the pB[PUbnlsEGFP] vector using methods described in the study by Manier *et al.* (46). These transgenic flies carry the *Ub-GFP* marker and a sperm-specific GFP marker (protamine-GFP, (*46*)). We backcrossed the *Ub-GFP* into *Dah* genetic background for 5 generations prior to the experiment. To produce males that had their insulin signaling pathway downregulated, we used *UAS-rpr* > *InsP3GAL* males. These males have their median neurosecretory cells ablated late in development by expressing *UAS-reaper* (*UAS-rpr*) under the control of the *InsP3-GAL4* (*InsP3GAL*) driver (*47*). This was achieved by crossing *InsP3GAL* males with *UAS-rpr* females, and collecting the male progeny (*UAS-rpr* > *InsP3GAL*). We backcrossed the *UAS-rpr* and *InsP3GAL* lines into the *whiteDahomey* genetic background for at least 5 generations prior to the experiment. We also used two sets of controls: The first control *InsP3GAL* were the sons of *InsP3GAL* males and *whiteDahomey* females and controlled for the effects of the GAL4 driver (*InsP3GAL*/+). The second control *UAS-rpr* were the sons of *whiteDahomey* males and *UAS-rpr* females and controlled for the effects of the UAS insert (*UAS-rpr*/+). Females used in the experiments were either *Dahomey* (in male ageing vials) or *spa* (in phenotypic assays). Males used in sperm competition assays were also *spa*. A sparkling eye (*spa*) stock was created by serially backcrossing *spa* into *Dah* background.

**S1.2 Experimental Design**

Upon eclosion, experimental virgin males were housed in groups of 12, either all males (Umated, “U”) or consisting of three virgin males and nine virgin *Dah* females (Frequently mated, “F”) (*19*). In the experiment with spermless males the F vials had one experimental *son-of-tudor* male, two *spa* males and nine virgin *Dah* females; the fertile *spa* males were provided to ensure that females were fertilised, to provide consistency with other experiments. Males were aged in their group vials for up to five weeks. Males from three age classes were used: 1 week (1w), 3 weeks (3w) and 5 weeks (5w) old (only 1w and 5w age classes were used in the spermless, insulin mutant and sperm viability experiments). The U flies were transferred once per week, and the F flies were transferred twice a week to fresh vials using light CO2 anesthesia at each transfer. During the transfers, dead or escaped females were replaced with similarly aged females. To minimize female co-ageing effects in the 5w (old) F group, females were replaced at 3 weeks with virgin 3-5 days old females, reared using the same procedures as above. To minimize density effects on mating opportunity in the F vials, two vials of the same treatment were merged when a single male was left in a vial owing to previous mortality or censoring. The males from F were merged into single sex groups of 10-12 males four/five days before assaying, in order to provide a consistent period of sexual rest prior to the assay point.

***The effect of aging and mating on reproductive traits and the seminal fluid proteins***

The day before the mating assays, 3-4 day old virgin *spa* females were placed individually in vials. On the day of mating assays, approximately 35 experimental *Dah* males from each ageing or mating treatment were added to the individually housed female vials and were given five hours to mate. Matings and associated parameters (mating latency and mating success) were recorded. The mated females were allowed to lay eggs for two days and the emerging offspring were counted to measure male fertility and offspring production. Then the females were transferred into a new vial with two 6-7 day old virgin *spa* males and were allowed to remate for five hours. Matings and associated parameters (remating latency and remating success) were recorded. The remated females were allowed to lay eggs for an additional two days and the emerging offspring were phenotyped and counted to measure paternity share.

The newly mated males were flash frozen in liquid nitrogen 30 minutes after the start of the mating. Another approximately 35 males from each treatment were flash frozen in liquid nitrogen without being exposed to females. We repeated this experiment to produce four independent biological replicates. We thawed flash frozen males and dissected their accessory glands and ejaculatory duct on ice in PBS buffer (*19*). 19 reproductive glands from males of the same treatment and replicate were pooled in 25μl PBS buffer on ice and sent for label-free quantitative proteomics sample preparation.

***The contribution of the seminal fluid proteome to reproductive ageing***

The day before the mating assays, a large number of 3-day old virgin *spa* females were placed individually in vials. On the day of mating assays, approximately 70 experimental spermless *son-of-tudor* and fertile control males from each age and mating treatment were added to the individually housed female vials and were given four hours to mate. Matings and associated parameters (mating latency and mating success) were recorded. Half of the mated females and a number of 6-day old virgin females (as controls) were subsequently transferred to new vials with two 4-day old virgin *spa* males, and allowed to remate for five hours. Matings and associated parameters (remating latency and remating success) were recorded. The other half of the mated females and a number of 8-day old virgin females (as controls) were transferred to yeast pasted vials and allowed to lay eggs for one day. The following day we counted the number of eggs laid in each vial.

***The effect of aging and mating on accessory gland size and sperm storage in males and mated females***

The day before the mating assays, a large number of 3-day old virgin *spa* females were placed individually in vials. On the day of mating assays, approximately 45 experimental *Ub-GFP* males from each ageing and mating treatment were added to the individually housed female vials and were given five hours to mate. Matings and associated parameters (mating latency and mating success) were recorded. The mated males were flash frozen in liquid nitrogen 30 minutes after the start of the mating. Another approximately 45 experimental *Ub-GFP* males from each treatment were flash frozen in liquid nitrogen without being exposed to females. We thawed flash frozen males and dissected their accessory glands and seminal vesicles on ice in PBS buffer. We imaged and measured the size of each seminal vesicle and accessory gland using a microscope calibration slide and ImageJ (*48*). The mated females were flash frozen in liquid nitrogen 90 minutes after the start of the mating. We thawed flash frozen females and dissected their spermatheca and seminal receptacle on ice in PBS buffer. We imaged and counted the number of sperm in both tissues using a fluorescence microscope and ImageJ (*48*).

***The effect of aging and mating on sperm production***

Adult testes were dissected in phosphate-buffered saline (PBS) and fixed in 5% formaldehyde for 20 min at room temperature, washed in PBX (PBS + 0.1% Triton X-100) for 15–20 min, and stained with rhodamine-phalloidin (0.1μM, Sigma-Aldrich, P1951) and DAPI (10μM, Invitrogen D1306) for 20 min at room temperature. Images were captured using a Zeiss LSM510 Confocal or an Olympus IX-81 motorized inverted microscope with XM-10 monochrome camera. Individualization complexes were scored using a 20x objective (*29*).

***The effect of aging and mating on sperm viability***

In the first experiment, we tested sperm viability inside the seminal vesicles in young and old U and F males. Approximately 30 *Dah* males from each age and mating treatment were dissected in ice-cold PBS buffer. The seminal vesicles were dissected from the testes and transferred to a 2.5μl drop of PBS Buffer on a new microscope slide and punctured to release sperm. The sample was covered to prevent evaporation and left for five minutes. Then 1.25μl of LIVE/DEAD stain (Thermo) was added and sperm viability was scored (*27*). The stain causes live sperm to fluoresce green and dead sperm to fluoresce red. If a sperm was stained both green and red, they were scored as dead (*27*). For each slide, four regions were imaged using a fluorescence microscope under both green and red filters, giving four pairs of images per slide. The images were processed in ImageJ and sperm viability for each sample was calculated as the proportion of sperm that were live (*48*). The experiment was repeated one more time to have two independent biological replicates. The imaging was done over four and five days in each replicate respectively.

In the second experiment we tested the effect seminal fluid produced by young and old U and F males on sperm viability. Approximately 30 *Dah* males from each age and mating treatment were dissected in ice-cold PBS buffer. The accessory glands were dissected and transferred to a 2.5μl drop of PBS Buffer on a new microscope slide and punctured to release seminal fluid. Another 1w male (standard) was dissected and the seminal vesicles were transferred to the same slide as the accessory glands. The seminal vesicles were punctured to release sperm; the seminal fluid and the sperm were mixed briefly with a pin. The sample was covered to prevent evaporation and left for an hour to give time for viability differences between treatments to develop (*27*). Then 1.25μl of LIVE/DEAD stain was added and sperm viability was scored. The imaging was done as detailed above.

***The effect of a lifespan extending manipulation on reproductive ageing***

The day before the mating assays, a large number of 4-5 day old virgin *spa* females were placed individually in vials. On the day of mating assays, approximately 35 experimental males from each transgenic line (*UAS-rpr* > *InsP3GAL, InsP3GAL*/+, *UAS-rpr*/+) age and mating treatment were added to the individually housed female vials and were given five hours to mate. Matings and associated parameters (mating latency and mating success) were recorded. The mated females were allowed to lay eggs for two days and the emerging offspring were counted to measure male fertility and offspring production. Then the females were transferred into a new vial with two 6-7 day old virgin *spa* males and were allowed to remate for five hours. Matings and associated parameters (remating latency and remating success) were recorded. The remated females were allowed to lay eggs for an additional two days and the emerging offspring were phenotyped and counted to measure paternity share. This experiment was run over two days and some of the experimental 5w males and *spa* males that failed to mate in the first day were tested again on the second day, which we controlled for in the statistical analyses.

**S1.3 Label-free Quantitative Proteomics**

***Sample Preparation***

All samples described above were stored at -80°C until sample preparation for proteomic analysis. The samples were macerated with a clean pestle and washed with 25μl of Pierce RIPA Buffer. Then they were digested using the standard gel-aided sample preparation (GASP) protocol as described previously (*19*, *49*). In brief, samples were reduced with 50mM DTT for 10 to 20 minutes. Protein lysate was mixed with an equal volume of 40% acrylamide/Bis solution (37.5:1. National Diagnostics) and left at room temperature for 30 minutes to facilitate cysteine alkylation to propionamide. 5ul TEMED and 5ul 10% APS were added to trigger acrylamide polymerization. The resulting gel plug was shredded by centrifugation through a Spin-X filter insert without membrane (CLS9301, Sigma/Corning). Gel pieces were fixed in 40% ethanol /5% acetic acid before 2 successive rounds of buffer exchange with 1.5M Urea, 0.5M Thiourea and 50mM ammonium bicarbonate which were removed with acetonitrile. Immobilized proteins were digested with trypsin (Promega) overnight and peptides extracted with two rounds of acetonitrile replacements. Peptides were first dried before desalting using Sola SPE columns (Thermo) and resuspended in 2% ACN, 0.1 % FA buffer prior LC-MS/MS analysis.

***LC-MS/MS***

Peptide samples were analysed on a LC-MS/MS platform, a Q-Exactive HF mass spectrometer (Thermo). After peptide loading in 0.1% TFA in 2% ACN onto a trap column (PepMAP C18, 300μm x5mm, 5μm particle, Thermo), peptides were separated on an easy spray column (PepMAP C18, 75μm x 500mm, 2μm particle, Thermo) with a gradient 2% ACN to 35% ACN in 0.1% formic acid in 5% DMSO. MS spectra were acquired in profile mode with a resolution of 60,000 with an ion target of 3x10^6^. The instrument was set to pick the 12 most intense features for subsequent MS/MS analysis at a resolution of 30,000, a maximum acquisition time of 45ms, an AGC target of 1x10^5^, an isolation width of 1.2Th and a dynamic exclusion of 27 seconds.

***Processing of MS Data***

LC-MS/MS data have been deposited to the ProteomeXchange Consortium (http:// proteomecentral.proteomexchange.org) via the PRIDE (*50*) partner repository with the data set identifier PXD009451. RAW files were imported into Progenesis QIP (version 3.0.6039.34628) using default settings. MS/MS spectra were exported as MGF files using the 200 most intense peaks without deconvolution for searching. The *Drosophila melanogaster* UniProt reference proteome was used as a search target in all cases. Database retrieval date was 30/03/2015 (21361 sequences). The dataset was searched in PEAKS using the following parameters: 10 ppm precursor mass accuracy, 0.05 Da fragment mass accuracy, Oxidation (M), Deamidation (N, Q) and Propionamide (K) as variable modifications, Propionamide (C) as a fixed modification, and two missed cleavage sites. We applied 1% FDR at peptide level (the search engine uses a target-decoy method for FDR estimation) and an additional Mascot ion score cutoff of 20 before importing search results into Progenesis, where protein quantification was calculated using the Top3 method. Quantitative protein data were further normalized/processed as described below.

**S1.4 Western Blot Assays**

Single males were ground in 10μl of sample buffer with a plastic pestle in 0.6 ml Eppendorf tubes. The samples were boiled for 5 min and spun for 2 min at 15000 rpm at RT. They were loaded into 13X13cm 5-15% gradient polyacrylamide (Amresco cat#M157) SDS gels with a 4% polyacrylamide stacking SDS gel. Gels were run at room temperature for 30 min at 110 volts, then moved to a 4^o^C room and run for 5 hours at 150 volts, until the dye front was approximately 9cm from the stacking gel. The gels were wet transferred to PVDF (Millipore cat# IPFL00010) membrane overnight at 4^o^C and 40 volts. Membranes were dried for at least 30 min to cross-link protein, then re-wet with 100% MeOH, and blocked with 5% milk in 1X TBST (0.1% Tween 20) for 1 hour at room temperature. Primary antibodies were diluted in 1% milk in 1X TBST (0.1% Tween 20) for at least 2 hours at room temperature, or overnight at 4^o^C. Membranes were rinsed 2X then washed 4X for 10 min each with 1X TBST (0.1% Tween 20). Secondary antibody was Goat Anti-Rabbit IgG (H+L) HRP (Jackson Immuno Research cat#111-035-003). Membranes were incubated for 1 hour in secondary antibody diluted 1:2000 in 5% milk in 1X TBST (0.1% Tween 20) for 1 hour at room temperature. Rinses and washes were repeated as above. Membranes were detected with Pierce ECL2 from Fisher cat#PI80196) and developed for 5 min at room temperature, then chemiluminescence was measured on a Typhoon scanner. Membranes were stripped with BME stripping buffer for 50 min at 50°C with shaking, then rinsed 3 times for 5 min each with 1XTBS, before blocking (as above) and adding an additional primary antibody. Primary antibody dilutions were 1:500 for 11864 Semp1, 1:5000 for Acp26Aa, 1:30000 for Acp36DE, 1:2000 for Acp62F, 1:1000 for CG9997, and 1:5000 for Sex peptide.

**S1.5 Statistical Analyses**

Data were analysed using RStudio 1.1.383 (*51*).

***The effect of aging and mating on reproductive traits and the seminal fluid proteins***

The proportion of survivors was compared between treatments using generalized linear models (GLMs) with a binomial error distribution corrected for overdispersion. The mating latency and remating latency of females were analysed using the *survival* package and a Weibull distribution. The proportion of infertile males (those producing 0 offspring) was tested using GLMs with a binomial error distribution. The number of offspring was analysed for fertile matings (i.e. those that produced at least 1 offspring), and was tested using GLMs with Poisson error distribution corrected for overdispersion. Paternity share was analysed using GLMs with a binomial error distribution corrected for overdispersion. The initial model included male age, mating history, their interaction and replicate number as fixed effects. Model selection was performed by backward stepwise elimination; non-significant (p>0.05) variables were eliminated from the model to arrive at the minimal adequate model. However replicate number was kept in the minimal model to control for this variation.

For the label-free quantitative proteomic dataset, only proteins identified with at least two unique peptides were included in the final dataset. From the 48 samples where 19 male reproductive tracts were pooled, we found a total of 1811 proteins, 1333 of which were identified by at least two unique peptides. We detected 117 known Sfps (*19*). 13 of these were ejaculatory duct specific. Quantitative data generated by Progenesis was normalised by log transforming the intensities [log2(x + 1)]. We followed the method of Keilhauer *et al.* (*52*) to determine a ‘background proteome’ for median centring purposes. Briefly, we calculated the standard deviation of the intensity profile for each identified protein, ranked the proteins according to the standard deviation of their profile, and selected the bottom 90% of the data. This ‘background proteome’ was used to median centre the distribution of each sample.

We focused our analyses on previously identified Sfps (*19*, *20*). Age and mating-related Sfp abundance differences, and transferred Sfp abundance differences, were analysed for each Sfp separately using linear mixed effect models. Here, the initial model included male age, mating history and their interaction as fixed effects, and replicate number as a random effect. Model selection was performed by backward stepwise elimination and the resulting p-values were corrected for multiple testing using Benjamini–Hochberg procedure. Age and mating-related Sfp abundance differences, and transferred Sfp abundance differences, were also analysed for all of the Sfps together using linear mixed effect models. Here, the initial model included male age, mating history, and their interaction, as fixed effects, and protein name and replicate number as random effects. Model selection was performed by backward stepwise elimination. We inferred the abundance of Sfps transferred to the female by subtracting the Sfp abundance of newly mated males from males not exposed to females within the same treatment and replicate. The heatmaps were made using the *pheatmap* package, and the data was mean centred (standardised) for each protein for better visualization. The lineplots were made using the *ggplot2* package. Age and mating-related compositional changes in the seminal fluid proteome and the transferred seminal fluid proteome were assessed using Principal component analyses (PCA) and linear mixed effect models. Again, the initial model included male age, mating history, and their interaction, as fixed effects, and replicate number as a random effect. Model selection was performed by backward stepwise elimination.

***The contribution of the seminal fluid proteome to reproductive ageing***

The number of eggs laid in one day following mating with a spermless male was analysed using two GLMs. The first one modelled the presence/absence of non-zero values using a binomial error distribution. The second one modeled the non-zero count data using a Poisson error distribution corrected for overdispersion. The remating latency of females immediately after mating to a spermless male were analysed using a Weibull distribution. The initial model included male age, mating history and their interaction as fixed effects. Model selection was performed by backward stepwise elimination. We also analysed the data including fertile control male and virgin female treatments. Here the number of eggs was analysed using a Poisson error distribution corrected for overdispersion. The remating latency of females were analysed using a Weibull distribution. The data was analysed separately for U and F groups. The initial model included treatment as a fixed effect.

***The effect of aging and mating on accessory gland size, sperm production, and sperm storage in males and mated females***

The size of accessory glands and seminal vesicles were analysed using a Gaussian distribution with an identity link function. The average number of cysts per testis was analysed using GLMs with Poisson error distribution corrected for overdispersion. The number of sperm in female sperm storage organs was analysed using two GLMs. The first one modelled the presence/absence of non-zero values using a binomial error distribution. The second one modeled the non-zero count data using a Poisson error distribution corrected for overdispersion. In each analysis the initial model included male age, mating history and their interaction as fixed effects. Model selection was performed by backward stepwise elimination.

***The effect of aging and mating on sperm viability***

Sperm viability inside the seminal vesicles was analysed using two GLMs. The first one modelled the presence/absence of any sperm using a binomial error distribution. The second one modeled the percentage data (percentage of live sperm within a sample) using a Poisson error distribution corrected for overdispersion. For both analyses the initial model included male age, mating history, their interaction, replicate number and day of experiment as fixed effects. Model selection was performed by backward stepwise elimination. However replicate number and day of experiment were kept in the minimal model to control for the variation introduced by these factors.

Sperm viability following mixing with seminal fluid from a different male was analysed using GLMs with a binomial error distribution corrected for overdispersion. The initial model included male age, mating, history, their interaction, replicate number and day of experiment as fixed effects. Model selection was performed by backward stepwise elimination. However replicate number and day of experiment were kept in the minimal model, as we wanted to control for the variation introduced by these factors.

***The effect of a lifespan extending manipulation on reproductive ageing***

The proportion of survivors at the time of assay was compared between treatments using GLMs with a binomial error distribution corrected for overdispersion. The mating latency and remating latency of females were analysed using a Weibull distribution. The proportion of infertile matings was analysed using GLMs with a binomial error distribution. The number of offspring was analysed for fertile matings using GLMs, with Poisson error distribution corrected for overdispersion. Paternity share from first mating was analysed using GLMs with a binomial error distribution corrected for overdispersion. The data was analysed separately for U and F groups. The initial model included male age, male line (*UAS-rpr* > *InsP3GAL, InsP3GAL*/+, *UAS-rpr*/+), their interaction, replicate number, day, and whether the vial had recycled males. Model selection was performed by backward stepwise elimination. However replicate number, day and whether the male was recycled were kept in the minimal model to control for these factors. Out of the 24 models we ran, in only four the two controls (*InsP3GAL*/+, *UAS-rpr*/+) had significantly different responses (Table S3). We therefore merged the two control genotypes as a single control to simplify subsequent analyses.

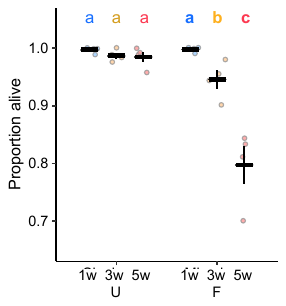

**Fig. S1: Old-F males are less likely to survive the experiment.** Age: χ ^2^_2_= 120.36; p< 0.0001; mating: χ ^2^_1_= 92.732; p< 0.0001; age and mating interaction: χ ^2^_2_= 6.246; p= 0.170 (n= 384 - 444, pooled from four replicates). Results are shown as means ± SEM. “U” stands for unmated and “F” stands for frequently mated males. Differences at p < 0.05 within mating groups and age categories are represented as different letters.

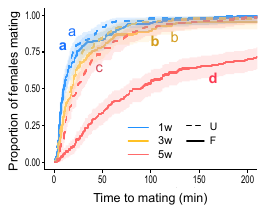

**Fig. S2: Old males show a significant reduction in copulation probability.** Age and mating interaction: χ ^2^_2_= 55.278; p < 0.0001 (n= 139-149). “U” stands for unmated and “F” stands for frequently mated males. Shaded areas are confidence intervals at 0.15 level. Differences at p < 0.05 within mating groups and age categories are represented as different letters.

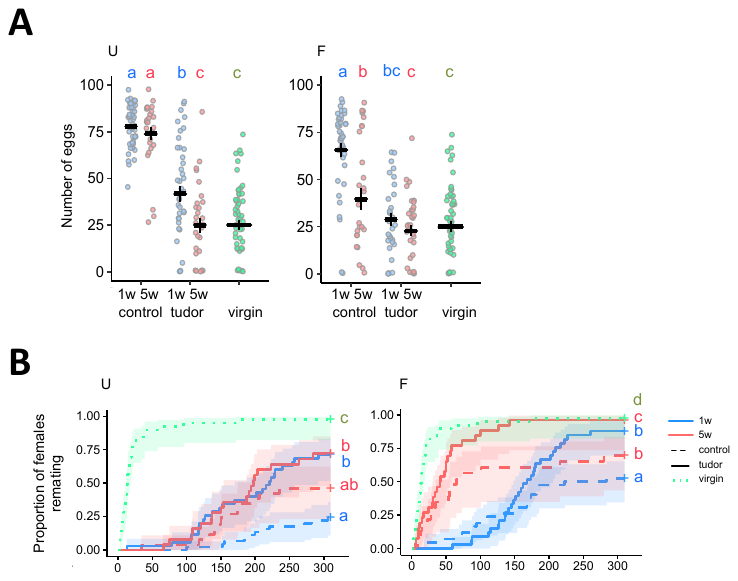

**Fig. S3: Old, spermless (*son-of-tudor*) males are poorer at stimulating female fecundity and suppressing female remating. (A)** Female fecundity (*U:* treatment: χ ^2^_4_= 2398.7; p< 0.0001; n=29-56) (*F:* treatment: χ ^2^_4_= 1405.9; p< 0.0001; n=29-56). **(B)** Female latency to remate (*U:* treatment: χ ^2^_4_= 137.923; p< 0.0001; n=25-45) (*F:* treatment: χ ^2^_4_= 99.421; p< 0.0001; n=23-42). “U” stands for unmated and “F” stands for frequently mated males. Shaded areas are confidence intervals at 0.15 level. Differences at p < 0.05 between treatments are represented as different letters.

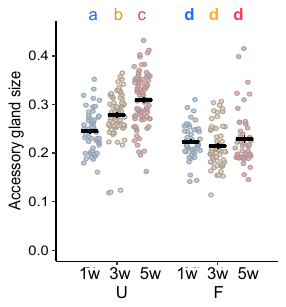

**Fig. S4: Aging and frequent mating impacts on accessory gland area (mm^2^).** Age and mating interaction: F^2^_2_= 10.685; p< 0.0001 (n= 49 - 84). “U” stands for unmated and “F” stands for frequently mated males. Results are shown as means ± SEM. Differences at p < 0.05 within mating groups and age categories are represented as different letters.

**Fig. S5:** **The gel mobility of a number of functionally important Sfps (Acp62F, Acp26Aa, Semp1, Acp36DE, Sex peptide, and CG9997) as determined by Western blots, in 1w and 5w males from U and F groups.** The abundance of each protein is predicted from the proteomic data and illustrated as a heatmap. Each lane is an individual male. “U” stands for unmated and “F” stands for frequently mated males.

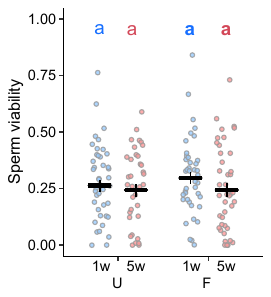

**Fig. S6**: **No evidence that sperm viability 1 hour after removal from the seminal vesicles responds to the age and mating status of the male contributing the seminal fluid.** Age: χ ^2^_1_= 33.132; p=0.226; mating: χ ^2^_1_= 2.562; p= 0.737; age and mating interaction: χ ^2^_1_= 11.579; p= 0.475 (n= 38 - 45). “U” stands for unmated and “F” stands for frequently mated males. Results are shown as means ± SEM. Differences at p < 0.05 within mating groups and age categories are represented as different letters.

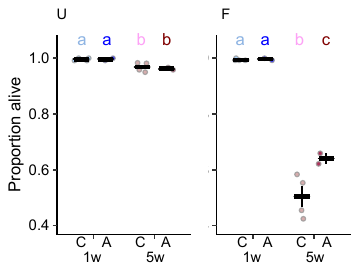

**Fig. S7: Old-F ablated males (*UAS-rpr* > *InsP3GAL*) are more likely to survive the experiment compared to Old-F control males (*InsP3GAL*/+ and *UAS-rpr*/+).** *U:* age: χ ^2^_1_= 19.288; p< 0.0001; line: χ ^2^_1_= 0.124; p= 0.692; age and line interaction: χ ^2^_1_= 0.015; p= 0.895. *F:* age: χ ^2^_1_= 556.52; p< 0.0001; line: χ ^2^_1_= 12.648; p= 0.005; age and line interaction: χ ^2^_1_= 0.013; p= 0.932 (n= 228 - 528, pooled from two replicates). “C” stands for control and “A” stands for ablated lines. “U” stands for unmated and “F” stands for frequently mated males. Results are shown as means ± SEM. Differences at p < 0.05 within lines and age categories are represented as different letters.

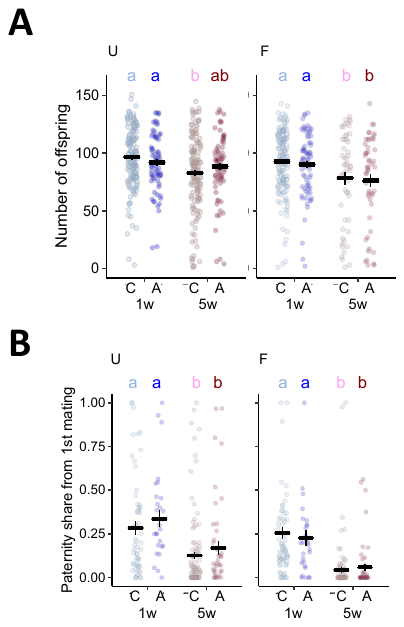

**Fig. S8: No evidence of differential offspring production and paternity share between ablated (*UAS-rpr* > *InsP3GAL*) and control (*InsP3GAL*/+ and *UAS-rpr*/+) males as a response to age and mating status.** **(A)** Offspring production (*U:* age: χ ^2^_1_=133.4; p< 0.0001; line: χ ^2^_1_=0.028; p= 0.953; age and line interaction: χ ^2^_1_= 26.518; p= 0.0681) (*F:* age: χ ^2^_1_=297.53; p< 0.0001; line: χ ^2^_1_=0.102; p= 0.923; age and line interaction: χ ^2^_1_=3.042; p= 0.6) (n= 51 – 153 pooled from three replicates). **(B)** Paternity share (*U:* age: χ ^2^_1_= 681.12; p< 0.0001; line: χ ^2^_1_= 2.582; p= 0.77; age and line interaction: χ ^2^_1_= 39.15; p= 0.256) (*F:* age: χ ^2^_1_= 1792; p< 0.0001; line: χ ^2^_1_= 0.008; p= 0.986; age and line interaction: χ ^2^_1_= 22.254; p= 0.365) (n= 30 – 112 pooled from three replicates). “C” stands for control and “A” stands for ablated lines. “U” stands for unmated and “F” stands for frequently mated males. Results are shown as means ± SEM. Differences at p < 0.05 within fly lines and age categories are represented as different letters.

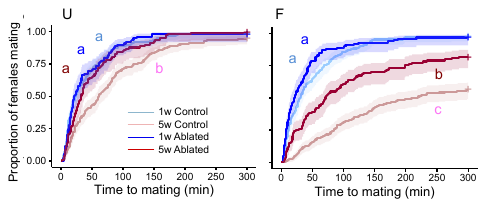

**Fig. S9: Females are more likely to mate with Old-F ablated males (*UAS-rpr* > *InsP3GAL*) compared to Old-F control males (*InsP3GAL*/+ and *UAS-rpr*/+).** *U:* age and line interaction: χ ^2^_1_= 7.447; p= 0.006 (n= 98-213). *F:* age and line interaction: χ ^2^_1_= 5.472; p= 0.019 (n= 100-227). “U” stands for unmated and “F” stands for frequently mated males. Shaded areas are confidence intervals at 0.15 level. Differences at p < 0.05 within mating groups and age categories are represented as different letters.

| ***Protein*** | ***Age qval*** | ***Mating qval*** | ***Age x Mating qval*** | ***Functional Category*** |
| --- | --- | --- | --- | --- |
| CG3097 | 0.001939 | 0.000001 | 0.0007 | Protease |
| CG10587 | 0.000328 | 0.000005 | 0.0010 | Protease |
| regucalcin | 0.002151 | 0.000001 | 0.0010 | Calcium ion binding |
| CG31413 | 0.006683 | 0.000010 | 0.0014 | Cell redox homeostasis |
| Sfp65A | 0.000127 | 0.000006 | 0.0014 | Unknown function |
| mfas | 0.000000 | 0.000002 | 0.0015 | Cell adhesion |
| Spn42Dd | 0.000267 | 0.000001 | 0.0015 | Protease inhibitor |
| CG31659 | 0.000025 | 0.000001 | 0.0028 | Lipid metabolism |
| CG34002 | 0.000215 | 0.000024 | 0.0029 | Unknown function |
| CG17093 | 0.005077 | 0.000004 | 0.0031 | Lipid metabolism |
| CG17097 | 0.007977 | 0.000009 | 0.0031 | Lipid metabolism |
| CG34051 | 0.000731 | 0.003108 | 0.0031 | Unknown function |
| Semp1 | 0.003534 | 0.001941 | 0.0031 | Protease |
| CG31883 | 0.215710 | 0.000001 | 0.0040 | Unknown function |
| Sems | 0.002151 | 0.000020 | 0.0040 | Post-mating behaviour |
| SP | 0.000328 | 0.000001 | 0.0040 | Post-mating behaviour |
| lectin-46Ca | 0.002152 | 0.000123 | 0.0045 | Post-mating behaviour |
| Sfp24F | 0.001384 | 0.000143 | 0.0048 | Carbohydrate interactions |
| CG34130-RA | 0.000122 | 0.000001 | 0.0060 | Protease |
| Acp26Ab | 0.001849 | 0.000076 | 0.0098 | Post-mating behaviour |
| CG31418 | 0.004048 | 0.000004 | 0.0098 | Unknown function |
| CG34129 | 0.008755 | 0.000010 | 0.0098 | Protease |
| CG3640 | 0.000267 | 0.000008 | 0.0098 | Unknown function |
| CG11608 | 0.032160 | 0.001008 | 0.0114 | Lipid metabolism |
| Acp26Aa | 0.012934 | 0.000495 | 0.0122 | Post-mating behaviour |
| CG10284 | 0.000132 | 0.000008 | 0.0122 | Defense/immunity |
| Acp53Ea | 0.000328 | 0.000048 | 0.0127 | Post-mating behaviour |
| lectin-29Ca | 0.001661 | 0.000003 | 0.0129 | Carbohydrate interactions |
| Spn28F | 0.007086 | 0.000012 | 0.0129 | Protease inhibitor |
| CG31419 | 0.001849 | 0.000004 | 0.0135 | Unknown function |
| CG31680 | 0.000063 | 0.000003 | 0.0176 | Unknown function |
| aqrs | 0.001384 | 0.000002 | 0.0188 | Post-mating behaviour |
| Obp56i | 0.000671 | 0.000000 | 0.0214 | Odorant binding |
| NUCB1 | 0.000024 | 0.000819 | 0.0220 | Defense/immunity |
| CG17472 | 0.000711 | 0.025065 | 0.0251 | Unknown function |
| CG17843 | 0.000671 | 0.000006 | 0.0259 | Cell redox homeostasis |
| Sfp38D | 0.001849 | 0.000008 | 0.0279 | Unknown function |
| CG17575 | 0.574146 | 0.042919 | 0.0328 | Post-mating behaviour |
| CG17919 | 0.025211 | 0.011913 | 0.0328 | Defense/immunity |
| CG9997 | 0.000267 | 0.000029 | 0.0344 | Post-mating behaviour |
| CG4847 | 0.014889 | 0.000161 | 0.0539 | Protease |
| CG9168 | 0.011594 | 0.023971 | 0.0539 | Catalytic activity |
| CG9519 | 0.011649 | 0.176250 | 0.0539 | Cell redox homeostasis |
| Est-6 | 0.024980 | 0.595285 | 0.0539 | Post-mating behaviour |
| CG10651 | 0.010840 | 0.011092 | 0.0565 | Unknown function |
| CG17242 | 0.003266 | 0.022477 | 0.0565 | Protease |
| Hexo2 | 0.001661 | 0.000002 | 0.0565 | Carbohydrate interactions |
| Spn38F | 0.079187 | 0.011565 | 0.0565 | Defense/immunity |
| Spn75F | 0.012124 | 0.004355 | 0.0583 | Protease inhibitor |
| CG10730 | 0.493557 | 0.083393 | 0.0587 | Catalytic activity |
| Acp36DE | 0.270410 | 0.000243 | 0.0655 | Post-mating behaviour |
| CG18284 | 0.000215 | 0.009889 | 0.0718 | Lipid metabolism |
| lectin-30A | 0.120793 | 0.000010 | 0.0718 | Carbohydrate interactions |
| Acp53C14a | 0.029885 | 0.000028 | 0.0725 | Unknown function |
| CG11598 | 0.294351 | 0.006846 | 0.0725 | Lipid metabolism |
| CG15116 | 0.015728 | 0.369419 | 0.0725 | Cell redox homeostasis |
| CG15117 | 0.014297 | 0.000005 | 0.0725 | Carbohydrate interactions |
| CG30395 | 0.855762 | 0.001899 | 0.0725 | Unknown function |
| CG31684 | 0.071129 | 0.001050 | 0.0725 | Lipid metabolism |
| lectin-46Cb | 0.002413 | 0.000737 | 0.0725 | Post-mating behaviour |
| Obp56g | 0.138601 | 0.017418 | 0.0747 | Odorant binding |
| CG1701 | 0.085978 | 0.054908 | 0.0821 | Unknown function |
| CG9029 | 0.047172 | 0.095215 | 0.0839 | Defense/immunity |
| betaTub85D | 0.645682 | 0.042919 | 0.0862 | DNA interactions |
| antr | 0.002805 | 0.000010 | 0.1017 | Post-mating behaviour |
| BG642312 | 0.372316 | 0.001851 | 0.1075 | Post-mating behaviour |
| Spn28B | 0.008040 | 0.000039 | 0.1075 | Protease inhibitor |
| Sfp78E | 0.077672 | 0.006576 | 0.1163 | Unknown function |
| CG14034 | 0.048468 | 0.000031 | 0.1209 | Lipid metabolism |
| CG2852 | 0.590456 | 0.004743 | 0.1209 | Catalytic activity |
| NLaz | 0.008446 | 0.023971 | 0.1379 | Lipid metabolism |
| Acp53C14c | 0.398788 | 0.000306 | 0.1536 | Unknown function |
| Sfp23F | 0.086745 | 0.002687 | 0.1536 | Protease inhibitor |
| Spn77Bb | 0.398788 | 0.776094 | 0.1548 | Protease inhibitor |
| Obp22a | 0.000731 | 0.000123 | 0.1622 | Odorant binding |
| Sfp26Ad | 0.016149 | 0.000474 | 0.1629 | Unknown function |
| Acp76A | 0.333139 | 0.069725 | 0.1715 | Protease inhibitor |
| CG6071 | 0.147006 | 0.685423 | 0.1715 | Protease |
| Ggt-1 | 0.138347 | 0.000160 | 0.1715 | Protease |
| CG6690 | 0.045501 | 0.000517 | 0.1849 | Cell redox homeostasis |
| Spn28Db | 0.148625 | 0.000006 | 0.1986 | Protease inhibitor |
| CG18067 | 0.000267 | 0.445143 | 0.2091 | Unknown function |
| Phm | 0.185761 | 0.095019 | 0.2143 | Catalytic activity |
| CG10041 | 0.924033 | 0.011473 | 0.2274 | Protease |
| CG10407 | 0.424137 | 0.007841 | 0.2274 | Unknown function |
| Acp29AB | 0.382685 | 0.000335 | 0.2289 | Post-mating behaviour |
| CG18135 | 0.415179 | 0.559060 | 0.2989 | Lipid metabolism |
| CG5162 | 0.012421 | 0.040758 | 0.3634 | Lipid metabolism |
| Sfp51E | 0.006425 | 0.000393 | 0.3674 | Unknown function |
| Sfp33A3 | 0.020628 | 0.004146 | 0.3882 | Catalytic activity inhibition |
| Obp56f | 0.097002 | 0.006955 | 0.4605 | Odorant binding |
| S-Lap7 | 0.187481 | 0.182767 | 0.5183 | Protease |
| CG15641 | 0.012124 | 0.000790 | 0.5300 | Unknown function |
| CG34034 | 0.859900 | 0.391358 | 0.5323 | Unknown function |
| Sfp24C1 | 0.014889 | 0.012834 | 0.5323 | Protease inhibitor |
| Mst57Dc | 0.031189 | 0.000158 | 0.5393 | Post-mating behaviour |
| BG642163 | 0.128093 | 0.012834 | 0.5524 | Unknown function |
| Obp51a | 0.464016 | 0.445143 | 0.5642 | Odorant binding |
| alphaTub84B | 0.001972 | 0.153894 | 0.6126 | DNA interactions |
| CG11112 | 0.070414 | 0.004466 | 0.6126 | Unknown function |
| Dup99B | 0.038606 | 0.332900 | 0.6126 | Post-mating behaviour |
| CG32833 | 0.016324 | 0.001256 | 0.6511 | Protease |
| Spn77Bc | 0.103226 | 0.026241 | 0.6905 | Protease inhibitor |
| Sfp70A4 | 0.294351 | 0.066354 | 0.6974 | Unknown function |
| CG11037 | 0.030346 | 0.000847 | 0.7095 | Protease |
| CG30486 | 0.294351 | 0.006955 | 0.7620 | Unknown function |
| CG31515 | 0.168643 | 0.022477 | 0.7620 | Protease inhibitor |
| Met75Ca | 0.487535 | 0.327324 | 0.7753 | Unknown function |
| Sfp35C | 0.110603 | 0.003618 | 0.8377 | Unknown function |
| Npc2b | 0.011594 | 0.332306 | 0.8645 | Hormone metabolism |
| Acp62F | 0.364605 | 0.004466 | 0.8903 | Post-mating behaviour |
| CG15635 | 0.590456 | 0.370730 | 0.9536 | Unknown function |
| Acp53C14b | 0.193640 | 0.019589 | 0.9736 | Unknown function |
| CG31704 | 0.528657 | 0.018633 | 0.9736 | Unknown function |
| CG34033 | 0.028763 | 0.000847 | 0.9736 | Unknown function |
| Obp56e | 0.814145 | 0.000635 | 0.9741 | Odorant binding |
| CG43145 | 0.291021 | 0.008410 | 0.9763 | Protease inhibitor |

**Table S1: The list of Sfps detected in this study and their functional categories.** The abundance of the top 40 Sfps show a significant differential response to age and mating after false discovery rate (FDR) correction (*Age x Mating qval*). Each Sfps individual response to age and mating after FDR correction are also included (*Age qval* and *Mating qval* respectively).

| ***Protein*** | ***Cluster*** | ***Age***  ***qval*** | ***Mating qval*** | ***Age x Mating qval*** | ***Functional***  ***Category*** |
| --- | --- | --- | --- | --- | --- |
| CG34034 | Yes | 0.860 | 0.391 | 0.532 | Unknown function |
| Obp51a | Yes | 0.464 | 0.445 | 0.564 | Odorant binding |
| Met75Ca | Yes | 0.488 | 0.327 | 0.775 | Unknown function |
| Spn77Bc | Yes | 0.103 | 0.026 | 0.691 | Protease inhibitor |
| Est-6 | Yes | 0.025 | 0.595 | 0.054 | Post-mating behaviour |
| Dup99B | Yes | 0.039 | 0.333 | 0.613 | Post-mating behaviour |
| CG5162 | Yes | 0.012 | 0.041 | 0.363 | Lipid metabolism |
| CG17242 | Yes | 0.003 | 0.022 | 0.057 | Protease |
| Obp56g | No | 0.139 | 0.017 | 0.075 | Odorant binding |
| CG18067 | No | 0.000 | 0.445 | 0.209 | Unknown function |
| Spn77Bb | No | 0.399 | 0.776 | 0.155 | Protease inhibitor |
| CG31704 | No | 0.529 | 0.019 | 0.974 | Unknown function |
| NLaz | No | 0.008 | 0.024 | 0.138 | Lipid metabolism |

**Table S2: The list of ejaculatory-duct specific Sfps detected in this study and their functional categories.** The top eight ejaculatory-duct specific Sfps clustered separately from the rest of the Sfps. Each Sfps individual response to age, mating and their interaction after FDR correction are also included (*Age qval*, *Mating qval* and *Age x Mating qval* respectively).

| ***Effect*** | ***Treatment*** | ***Estimate*** | ***Standard error*** | ***T value*** | ***P value*** |
| --- | --- | --- | --- | --- | --- |
| Infertile matings | 1wU | -0.874 | 0.761 | -1.147 | 0.251 |
|  | 5wU | -0.104 | 0.353 | -0.293 | 0.769 |
|  | 1wF | -1.066 | 0.891 | -1.196 | 0.232 |
|  | 5wF | 0.855 | 0.445 | 1.921 | 0.055 |
| Remating latency | 1wU | 0.475 | 0.228 | 2.080 | 0.037 |
|  | 5wU | 0.589 | 0.248 | 2.380 | 0.018 |
|  | 1wF | 0.176 | 0.181 | 0.970 | 0.333 |
|  | 5wF | -0.415 | 0.253 | -1.640 | 0.101 |
| Proportion alive | 1wU | -1.149E-15 | 3.890e-05 | 0 | 1 |
|  | 5wU | 0.993 | 0.149 | 6.650 | 0.095 |
|  | 1wF | -0.697 | 1.065 | -0.655 | 0.631 |
|  | 5wF | -0.551 | 0.090 | -6.107 | 0.103 |
| Offspring number | 1wU | -0.030 | 0.040 | -0.740 | 0.460 |
|  | 5wU | -0.079 | 0.064 | -1.238 | 0.218 |
|  | 1wF | -0.001 | 0.045 | -0.011 | 0.991 |
|  | 5wF | -0.286 | 0.160 | -1.785 | 0.081 |
| Paternity share | 1wU | 0.529 | 0.457 | 1.158 | 0.252 |
|  | 5wU | 0.201 | 0.368 | 0.546 | 0.586 |
|  | 1wF | -0.112 | 0.294 | -0.381 | 0.704 |
|  | 5wF | -2.660 | 1.151 | -2.312 | 0.023 |
| Mating latency | 1wU | -0.388 | 0.153 | -2.540 | 0.011 |
|  | 5wU | 0.224 | 0.136 | 1.640 | 0.101 |
|  | 1wF | 0.042 | 0.150 | 0.280 | 0.778 |
|  | 5wF | 0.005 | 0.173 | 0.030 | 0.976 |

**Table S3: Pairwise comparisons between the two control genotypes (*InsP3GAL*/+ and *UAS-rpr*/+) within each age and mating treatment.** These are 1w and 5w old U and F treatments. The response variables are the proportion of infertile matings, female latency to remating, the proportion of flies to survive the experiment, offspring production, paternity share of the first male and female latency to mating. “U” stands for unmated and “F” stands for frequently mated males. Differences at p < 0.05 are given in red.
